# Free energy perturbation calculations of mutation effects on SARS-CoV-2 RBD::ACE2 binding affinity

**DOI:** 10.1101/2022.08.01.502301

**Authors:** Alina P. Sergeeva, Phinikoula S. Katsamba, Jared M. Sampson, Fabiana Bahna, Seetha Mannepalli, Nicholas C. Morano, Lawrence Shapiro, Richard A. Friesner, Barry Honig

## Abstract

The strength of binding between human angiotensin converting enzyme 2 (ACE2) and the receptor binding domain (RBD) of viral spike protein plays a role in the transmissibility of the SARS-CoV-2 virus. In this study we focus on a subset of RBD mutations that have been frequently observed in infected individuals and probe binding affinity changes to ACE2 using surface plasmon resonance (SPR) measurements and free energy perturbation (FEP) calculations. Our SPR results are largely in accord with previous studies but discrepancies do arise due to differences in experimental methods and to protocol differences even when a single method is used. Overall, we find that FEP performance is superior to that of other computational approaches examined as determined by agreement with experiment and, in particular, by its ability to identify stabilizing mutations. Moreover, the calculations successfully predict the observed cooperative stabilization of binding by the Q498R N501Y double mutant present in Omicron variants and offer a physical explanation for the underlying mechanism. Overall, our results suggest that despite the significant computational cost, FEP calculations may offer an effective strategy to understand the effects of interfacial mutations on protein-protein binding affinities and in practical applications such as the optimization of neutralizing antibodies.

## Introduction

The ability to accurately predict binding affinity changes upon mutations of interfacial residues is a problem of significant importance, ranging from the general problem of understanding of interaction specificity and the design of therapeutics such as potent monoclonal antibodies that target antigens to revealing the mechanism of action of cancer driver mutations. Multiple approaches to the problem have been developed including machine learning methods (1–5), statistical potentials (6) and various force field related scoring functions (7–11) embedded in programs such as FoldX (11) and Rosetta (7). Each approach is associated with its own set of issues, such as conformational changes upon mutation, that are nicely discussed in reviews of Bonvin and co-workers (12). Moreover, some methods succeed on some test sets and fail on others, suggesting either over-training or simply that some protein-protein interfaces have different properties than others. Issues of experimental validation can also arise (13); not all experimental methods are equally accurate and, as discussed below, the nuances of the experimental system can have significant effects on the outcome.

Detailed atomic-level simulations have not been extensively applied to the prediction of mutation effects, in part due to the computational requirements involved. Free-energy perturbation (FEP) methods have the potential to impact the field as physics-based force fields are, in principle, agnostic to the system being studied. Most current applications have involved the optimization of ligand-protein interaction in the context of small molecule drug design (reviewed in (14)) but recent publications have begun to explore the use of FEP methods to the study of protein-protein interactions (PPIs); specifically, to the effects of interfacial mutations on protein-protein binding free energies (8, 9, 15–19). This is an inherently complex problem since, as opposed to relatively rigid ligand binding pockets, protein-protein interfaces are often quite large and less constrained so that they can more easily undergo conformational change as a result of a mutation. Moreover, FEP calculations involve a complex computational infrastructure and are extremely time consuming. However, fast graphical processing units (GPUs) made such calculations feasible and a number of recent publications, involving different software packages, suggest that the methodology has reached the point that good correlation with experiment is to be expected (8, 9, 15–19). Clearly, if FEP methods are capable of providing meaningful results, then in many applications, the computational cost will be worthwhile.

Here we explore the ability of FEP calculations to reproduce the effects of mutations on the binding of the receptor binding domain (RBD) of the SARS-CoV-2 spike protein with the human angiotensin converting enzyme 2 (ACE2) using the FEP+ implementation (see Methods). Given that the pathogen entry into the host cell is mediated by RBD::ACE2 binding, the problem has attracted considerable interest and multiple experimental (20–46) and computations studies (33, 47–63) have been reported. We chose to study a set of 23 frequently observed RBD mutations (Table S1) located in the RBD::ACE2 interface (Fig. 1) of Alpha, Beta, Gamma, Delta, or Omicron SARS-CoV-2 variants (Table S2). Surface Plasmon Resonance (SPR) experiments were carried out for each and compared to previous experimental work(20–46). A number of computational methods were applied to predict binding free energy changes upon mutations, ΔΔG. We found FEP to be the best performer and, in particular, provides a good starting point for an experimental program to optimize protein-protein binding affinities. In particular FEP trajectory analysis has allowed us to characterize the underlying biophysical effects that produce stabilizing mutations. In this regard, we show that FEP successfully recapitulates the stabilizing epistatic effect of the Q498R N501Y mutant present in every Omicron variant. The ability to anticipate non-additive effects of multiple mutations is likely to be an important element of future efforts in protein interface design.

**Figure 1.**
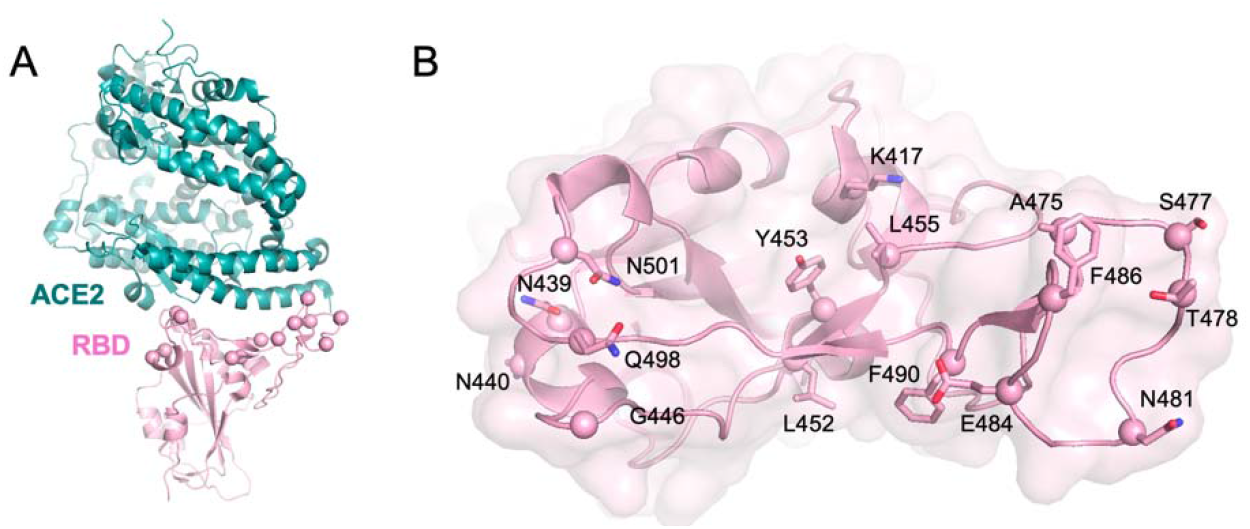
ACE2/RBD system of interest. (A) Ribbon representation of the ACE2/RBD complex. (B) Side chains of interfacial RBD residues in contact with ACE2 probed in this study are shown in stick representation with C⍰ atoms shown as spheres.

## Results

### SPR Measurements of Binding Affinity Changes

The second column in Table 1 lists experimental changes in binding affinity (ΔΔG) of the ACE2::RBD complex when RBD is mutated. Of 23 single point RBD mutations probed, only four were identified as stabilizing with ΔΔG values ≤ −0.4 which is our measure of experimental accuracy (see Methods): N501Y, Y453F, S477N and N501T (Table 1, see Fig. S1 for corresponding fitted SPR data and Methods for experimental detail). The third column lists ΔΔG values from the deep mutational scanning study of Starr et al.(20). Although the methods are quite different and our SPR results are obtained with monomeric ACE2 while Starr et al. used dimeric ACE2, the results are in good agreement (high Pearson correlation coefficient (PCC)=0.9 and low root mean square error (RMSE)=0.2 kcal/mol). Given that our calculations are carried out on a structure containing monomeric ACE2 and the likelihood that the SPR results are more accurate than the high-throughput yeast display values, we use the SPR values to compare to computational predictions.

**Table 1.** Experimental ACE2/RBD binding affinity changes for RBD mutants.

| Mutation | $\Delta\Delta G$<br>(SPR,<br>this study) | $\Delta\Delta G$<br>(Yeast Display,<br>Starr et al.)<sup>(20)</sup> | $\Delta\Delta G$ (other studies)<br>SPR<sup>(21-33)</sup><br>BLI<sup>(34-46)</sup><br>Yeast Display<sup>(64)</sup><br>(range) |
| --- | --- | --- | --- |
| N501Y | -0.8 | -0.3 | (-1.7 ... -0.5) |
| Y453F | -0.7 | -0.3 | (-1.2 ... -0.8) |
| S477N | -0.5 | -0.1 | (-0.6 ... -0.1) |
| N501T | -0.5 | -0.1 | (-0.9 ... 0.0) |
| N439K | -0.1 | 0.0 | (-0.4 ... 0.0) |
| N440K | 0.0 | -0.1 |  |
| F490S | 0.0 | 0.0 |  |
| L452M | 0.0 | -0.1 |  |
| L452R | 0.0 | 0.0 | -0.2 |
| E484Q | 0.1 | 0.0 |  |
| T478K | 0.1 | 0.0 | 0.2 |
| N481K | 0.1 | 0.0 |  |
| E484K | 0.1 | 0.1 | (-0.6 ... 0.3) |
| Q498R | 0.2 | 0.1 | (-0.5 ... 0.5) |
| S477I | 0.2 | 0.1 |  |
| G446V | 0.2 | 0.4 | 0.5 |
| T478R | 0.2 | 0.1 |  |
| S477R | 0.3 | 0.0 | -0.5 |
| A475V | 0.3 | 0.2 | 0.2 |
| L455F | 0.4 | 0.3 |  |
| K417T | 0.4 | 0.4 | (0.1 ... 0.7) |
| F486L | 0.6 | 0.6 | 0.9 |
| K417N | 0.6 | 0.6 | (-0.5 ... 1.1) |

Experimental binding affinity difference values were calculated based on binding affinity ( $K_D$ ) measurements for wild-type (WT) and single mutant (MT) proteins using the following formula:

Table 1 also lists ΔΔG values obtained previously with SPR (21–33) and other experimental methods; bio-layer interferometry (BLI) (34–46) and yeast display (64). Direct comparisons are difficult since, for example, different constructs were used in different experiments and different proteins were used as analytes in some SPR experiments (see Methods for details). The differences in constructs result from the choice of monomeric vs. multimeric forms of interacting proteins as well as the selection of protein domain boundaries. Nevertheless, overall, there is good agreement among most experimental results with the outliers attributable to the factors mentioned here. Moreover, there is good consensus regarding the identity of the most stabilizing mutations. For example, our results for N501Y (−0.8 kcal/mol), Y453F (−0.7 kcal/mol), S477N (−0.5 kcal/mol) and N501T (−0.5 kcal/mol) are in good agreement with previously published values (20–32, 34, 36–38, 40–46, 65) regardless of constructs/experimental setup differences: N501Y (−1.7 < ΔΔG < −0.5), Y453F (−1.2 < ΔΔG < −0.8), S477N (−0.6< ΔΔG < −0.1) and N501T (−0.9 < ΔΔG < 0.0).

### Computational Prediction of the Effect of Mutations on Binding Affinity

Table 2 presents FEP results for the 23 experimental ΔΔG values obtained from SPR measurements listed in Table 1. Correlation plots for the data in Table 2 are given in Figures 2 and S2. The overall performance of FEP is in line with previous work (8, 9, 14, 66–68); a PCC of 0.6 and an RMSE of 0.8 kcal/mol (Table 2). The FEP calculations clearly predict that all four stabilizing mutations, N501Y, N501T, S477N, and Y453F have ΔΔG values < 0 although the prediction for S477N is a weak one. Of note, previous FEP calculations on N501Y (53, 55, 56) have yielded results very similar to ours attesting to the robustness of the method. The most significant failure of FEP is its prediction that A475V is stabilizing when the SPR results indicate that it is weakly destabilizing. A likely explanation for this result is that A475 is located close to the N-terminal residue of ACE (Q18) for which no coordinates were assigned in the crystal structure (69). Another problematic result, a largely overestimated ΔΔG for Q498R, will be discussed below. The FEP results in Table 2 and Figure 2 were obtained from 100 ns simulations. Extensive prior work (8, 9) has demonstrated that shorter simulations are often inadequate for some mutations, typically those involving residues with a low degree of solvent exposure (i.e. partially or fully buried at the protein-protein interface). The results of 10 ns FEP simulations are shown in the Supplementary Material (Table S3 and Fig. S2) for comparison. Of particular note is the N501Y mutation, which is predicted to be destabilizing at 10 ns but is then correctly predicted to be stabilizing at 100 ns.

**Table 2.**
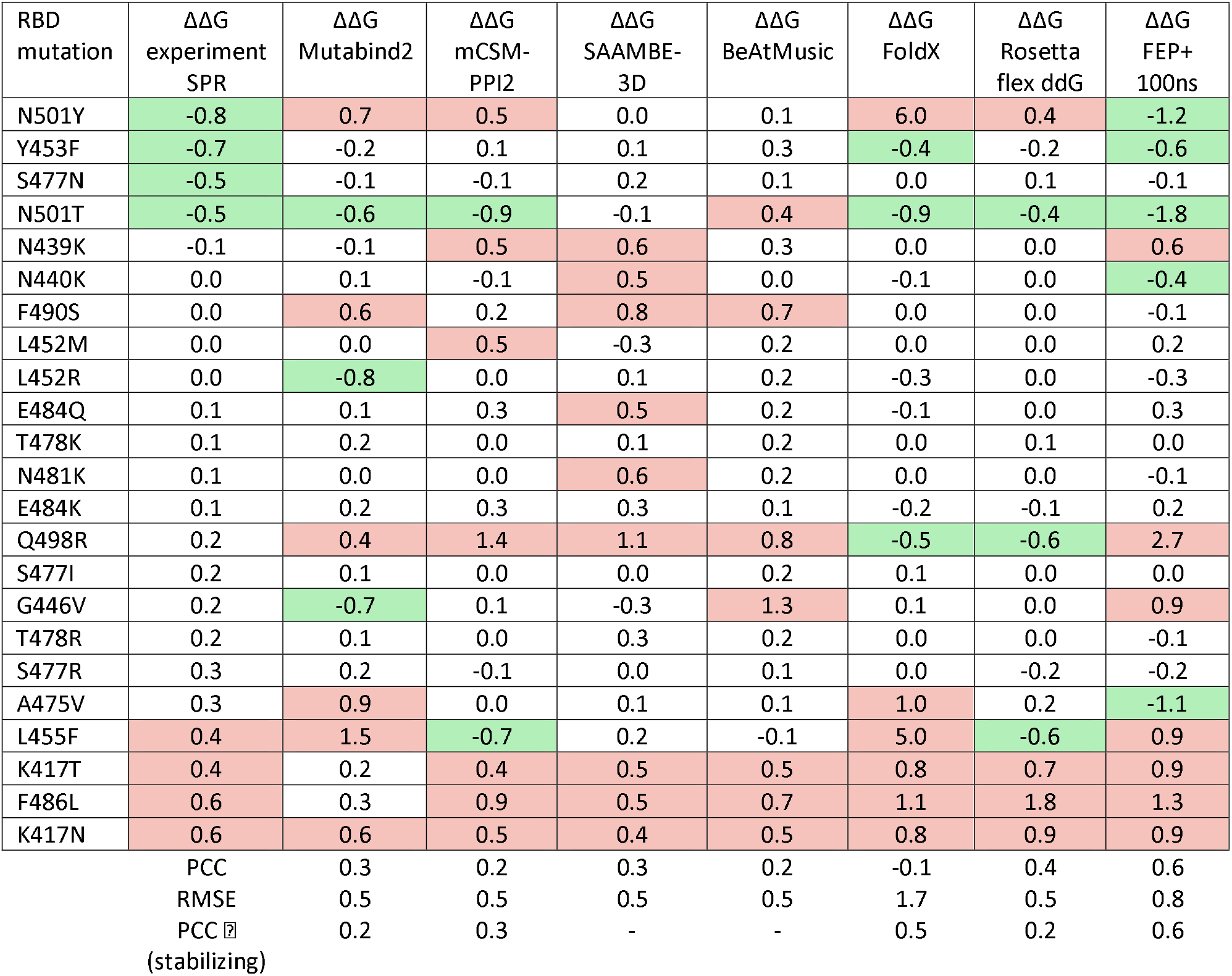
Calculated ACE2/RBD binding affinity changes for RBD mutants. Pearson correlation coefficient (PCC), Pearson phi for stabilizing mutations (PCC ⍰) and root mean square error (RMSE) are calculated for every method tested based on comparison to SPR results in Table 1. Stabilizing mutations with ΔΔG ≤ −0.4 kcal/mol in green, destabilizing mutations with ΔΔG ≥ 0.4 kcal/mol in red. Experimental binding affinity difference values were calculated based on SPR binding affinity (K_D_) measurements for wild-type (WT) and single mutant (MT) proteins using the following formula: ΔΔG = RT ln (K_D_(MT) / K_D_(WT)), in units of kcal/mol. Correlation plots for all theoretical methods are provided in Fig. S2. Calculations were performed on a crystal structure of ACE2/RBD (PDBID 6M0J). Protein specific residue numbering of all the mutants as in Uniprot ID P0DTC2.

**Figure 2.**
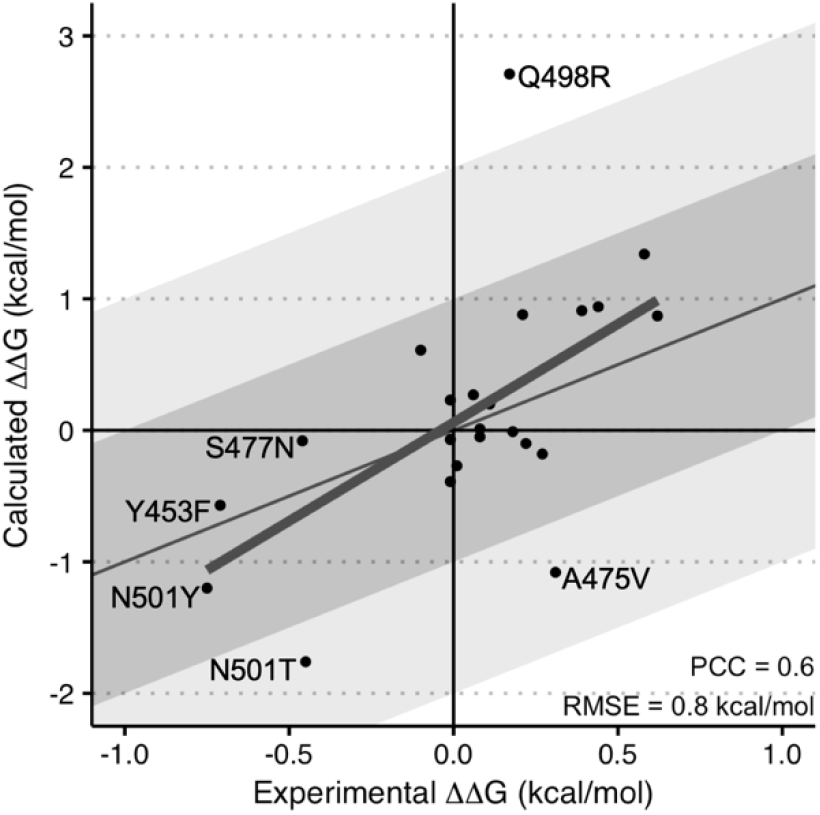
Correlation plot between FEP and SPR binding affinity changes. FEP results are calculated at 100ns. Mutations discussed in the text are labeled. The ΔΔG values are given in Table 2 along with comparison of FEP performance to other theoretical methods.

Machine learning (ML) methods; Mutabind2 (5), mCSM-PPI2 (3) and SAAMBE-3D (1) uniformly have a weak correlation with experiment (PCC<0.4). As we have pointed out previously (60) ML methods tend to overpredict destabilizing mutations presumably due to the preponderance of destabilizing mutations in training sets. A related factor likely accounts for relatively low RMSEs of ML methods since most mutations in training sets have only small effects on binding affinities. BeAtMuSiC evaluates mutation effects using a statistical potential (6) while FoldX uses an empirical physics-based force field (10, 11). Both methods assume a rigid backbone although FoldX allows for side chain rearrangement upon mutation. Neither method produces a meaningful correlation with experiment. BeAtMuSiC identifies no stabilizing mutations though FoldX correctly identifies N501T and Y453F. Of note, we have found FoldX to be quite effective in predicting mutation effects in two families of neuronal adhesion proteins that bind via a canonical interface not prone to backbone rearrangement (70).

Other than FEP, Rosetta flex ddG (7) is the only method that allows for backbone flexibility. Its PCC is still significantly less than that of FEP and, as can be seen in Table 2, it predicts two of the four stabilizing mutations (although the prediction for Y453F is a weak one). Nevertheless, even its partial success highlights the need to account for the ability of proteins to relax in response to interfacial mutations.

### Physical Insights from Trajectory Analysis

An important feature of the FEP approach is that analysis of trajectories can reveal insights that can be useful in the design of mutations with desired properties. The following cases offer examples of the type of information that can be extracted from the simulations.

The N501Y mutation is responsible for a high infectivity and transmissibility of the Alpha variant of SARS-CoV-2 (71) and has the largest stabilizing effect (Table 1). The N501T mutation found in the SARS-CoV-2 variants transmitted from mink to humans (72, 73) occurs at the same position. Analysis of FEP trajectories reveals that the stabilization effect associated with N501Y and N501T mutations is due to substitution of the asparagine with less polar side chains of tyrosine and threonine. N501 has only one of its polar groups satisfied in the wild-type (WT) structure while, throughout the course of the relevant trajectories, the hydroxyl groups of both tyrosine and threonine participate in hydrogen bonds (see dashed lines, Fig. 3A). In addition to enhanced stability due to the absence of unsatisfied hydrogen bonds, both mutants undergo stabilizing interactions with Y41 of ACE2; the aromatic ring of Y501 participates in *π*-*π* stacking interactions (see purple lines in Fig. 3A) while the methyl group of T501 forms a hydrophobic contact with the aromatic ring of Y41 (see gray shading in Fig. 3A). Of note, previous studies have identified the role of *π*-*π* stacking as a source of stabilization of the N501Y mutant (47, 53, 56, 58) but the role of the unsatisfied hydrogen bond in the WT protein has not been emphasized.

**Figure 3.**
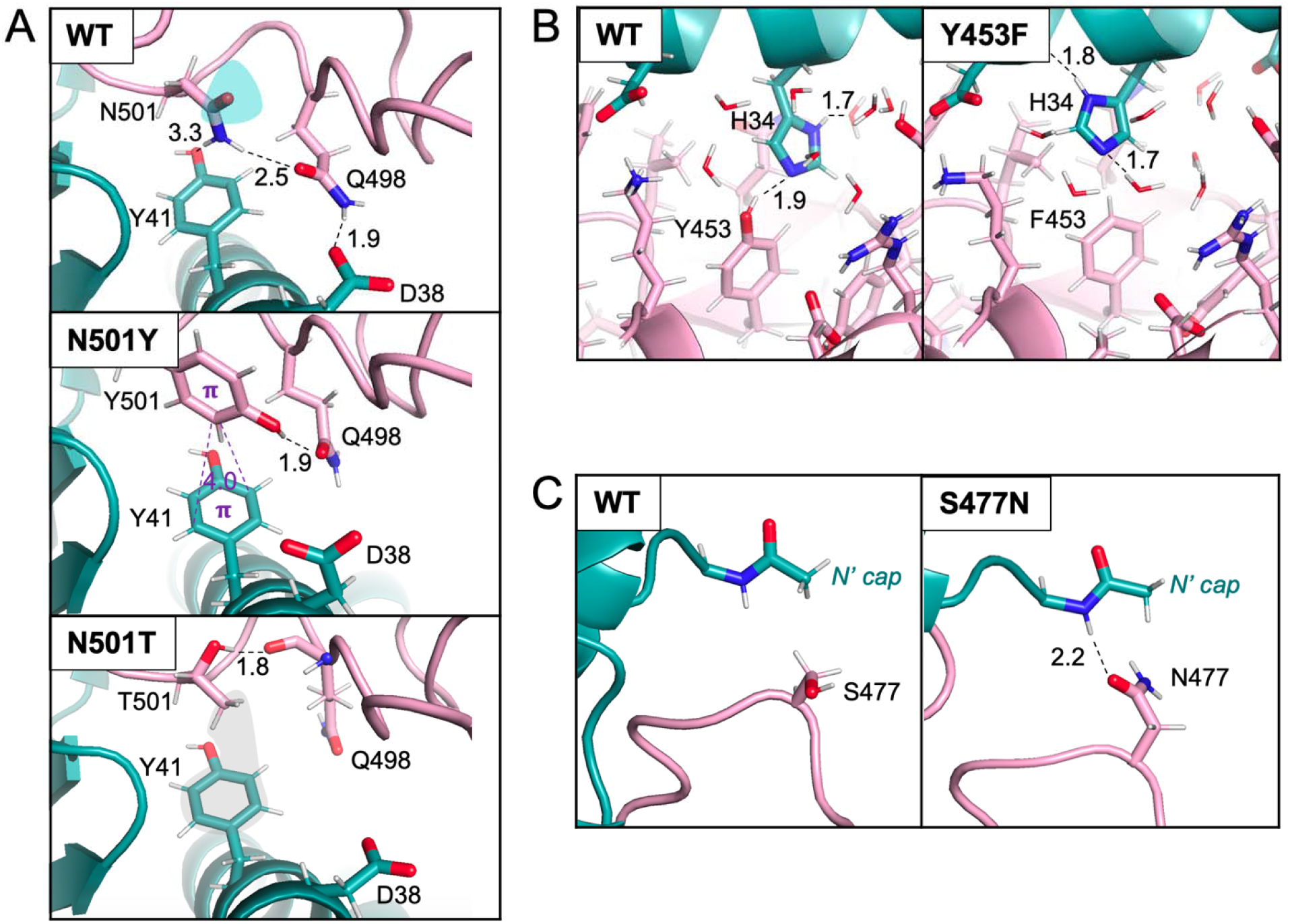
Structural origins of stabilizing ACE2/RBD interactions. Closeups of the ACE2(cyan)/RBD(pink) showing key interactions in wild-type (WT) and mutant proteins. Hydrophobic contacts are in grey, - interactions are in purple, hydrogen bonds are shown as black dashed lines, unsatisfied polar group of N501 is in cyan, favorable hydrophobic contact between T501 and Y41 in grey, and all distances are in Å.

Analysis of trajectories associated with the Y453F mutation shows that in the WT protein, a hydrogen bond between the hydroxyl group of Y453 and the Nε atom of H34 is present in ~25% of the WT trajectory (Fig. 3B) while, in most cases, these two residues form hydrogen bonds with trapped solvent molecules. In the Phe mutant, there is no need to satisfy the buried hydroxyl of the tyrosine while the Nε of H34 is satisfied by structured waters or backbone atoms (see dashed lines showing hydrogen bonds in Fig. 3B). Thus, the enhanced stability of the mutant is likely due to the greater hydrophobicity of a Phe relative to a Tyr. Of note, our trajectory analysis is in agreement with a previous study comparing crystallographic structures of the WT and Y453F mutant (32).

The simulations correctly predict that the S477N mutation is stabilizing but only weakly so. This residue faces the N-terminal of ACE2, and, specifically, the Q18 residue for which coordinates were missing in the crystal structure and, hence, modelled as an acetyl group cap in our simulations (Fig. 3C). Thus, the calculations may suffer from conformational uncertainties in this region. The trajectories reveal that the longer Asn side chain makes more contacts with the ACE2 N-terminal than does the WT Ser. We show only one snapshot from wild-type and mutant trajectories in Fig. 3C, but in reality, due to the flexibility of the ACE2 N-terminal, no specific contact is retained throughout the simulations.

The largest error in the FEP calculations is for the Q498R mutant whose destabilizing effect is over-predicted by about 2.5 kcal/mol. Analysis of the 100 ns Q498R trajectory reveals that the Arg side chain samples different conformations with the most prevalent state (~53%) involving an unfavorable polar-hydrophobic contact between N501 and the aliphatic chain of the Arg, leaving both polar groups of the Asn unsatisfied (Fig. 4A). This is likely a contributor to the destabilizing ΔΔG value. As we have noted previously (9), a computationally unfavorable outlier of this magnitude is typically attributable to the failure of the molecular dynamics trajectories (100 ns in this study) to achieve a converged reorganization of the protein structure. To test this possibility, we carried out a 300 ns simulation which appears to reach convergence at ~200ns (Fig. S3) where the destabilizing effect of Q498R is calculated to be 1.9 kcal/mol; reduced from 3.6 kcal/mol at 10 ns and 2.7 kcal/mol at 100 ns. It may well be the case that the system has converged to a metastable state at 300ns and that there are lower energy states not sampled in the course of the simulation.

**Figure 4.**
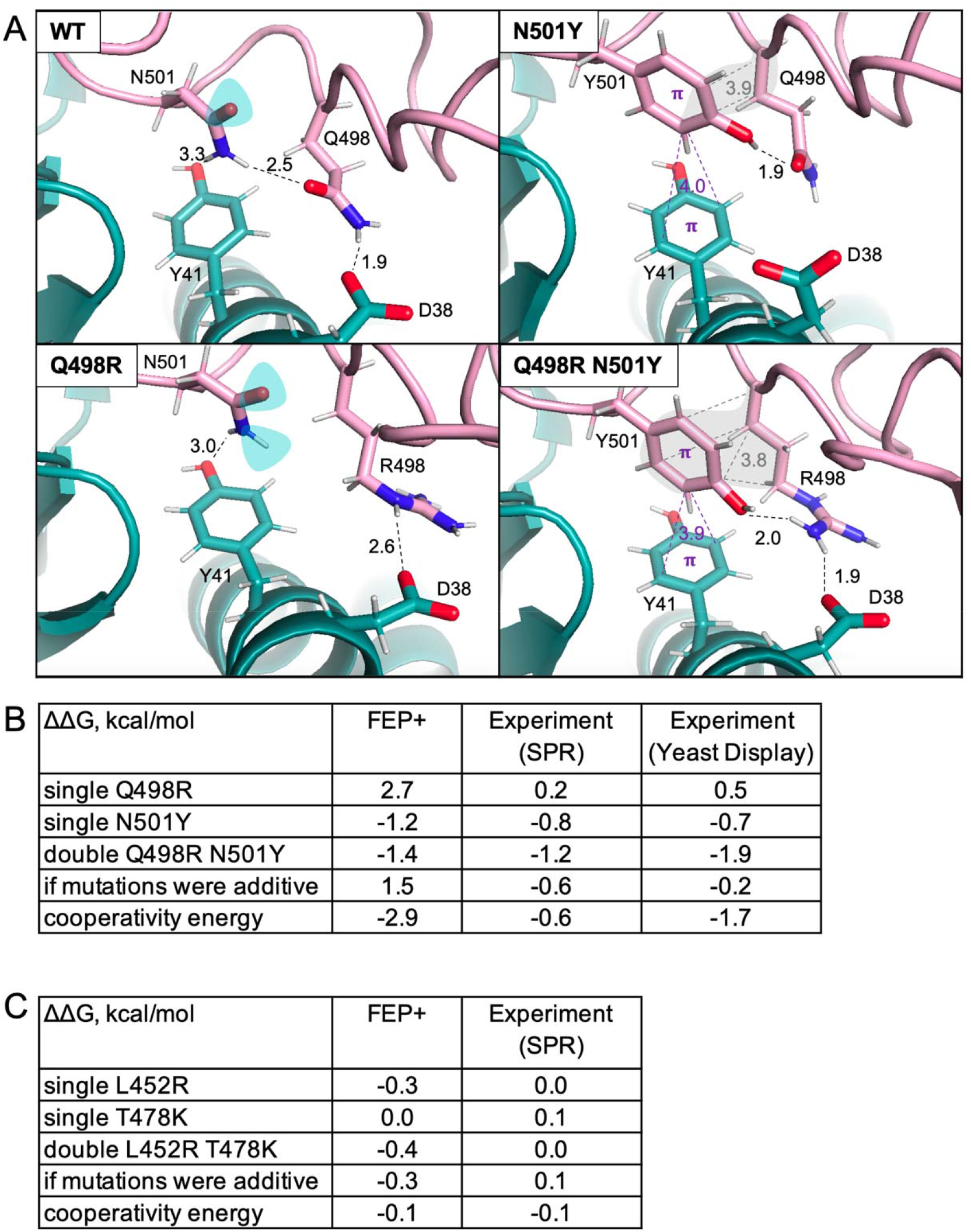
Epistatic effect of the double Q498R N501Y mutant. (A) A closeup of the complex between ACE2 (in cyan) and RBD (in pink) showing chemical interactions involving RBD residues 501 and 498. Hydrophobic contacts are in grey, - interactions are in purple, hydrogen bonds are shown as black dashed lines, unsatisfied polar groups are in cyan, and all distances are measured in Å. (B) Cooperativity of the Q498R N501Y double mutant probed by FEP+ and SPR computed as a difference between a hypothetical ΔΔG (if the double mutant had an additive effect of two single point mutations) and the actual ΔΔG of the double mutant. Yeast display assay values are from Zahradnik et a (64). (C) Absence of cooperativity probed by FEP+ and SPR for the L452R T478K double mutant. Experimental result for the Delta RBD variant is from Liu et al. (81) (see Methods for details), ΔΔG of single mutants are from the current study (Table 2). The FEP+ results are from 100ns trajectories.

### Epistatic Effect of the Q498R N501Y Double Mutant in the Omicron variant

Zahradnik et al. (64) demonstrated that the Q498R N501Y double mutant is more stabilizing than the additive effect of two single point mutations as estimated by their yeast display assay (a “cooperativity energy” of ~ −1.7 kcal/mol). Our SPR results on the single Q498R and N501Y mutants predict, if their effects were additive, that the double mutant would be stabilizing by −0.6 kcal/mol (−0.8 + 0.2 kcal/mol for the single mutations, respectively) while the experimental value for the double mutant is −1.2 kcal/mol yielding a cooperativity energy of −0.6 kcal/mol (see Fig. 4B for the experimental ΔΔG values and Fig. S1 for corresponding fitted data and dissociation constants). While the experimental SPR and yeast display ΔΔG values differ, both methods indicate that a substantial epistatic effect is playing a role, perhaps contributing to the greater infectivity of the Omicron variants where these mutations are present.

FEP calculations on the double mutant predict a ΔΔG of −1.4 kcal/mol whereas the predicted additive effect of the two single mutants (Table S3) is +1.5 kcal/mol – corresponding to a cooperativity energy of −2.9 kcal/mol (Fig. 4B). Thus, the 100 ns calculations successfully predict the existence of cooperativity. A detailed examination of the structure of the double mutant provides a compelling physical interpretation as to why it behaves so differently from the single Q498R mutation. In the double mutant, Arg 498 forms a favorable pairing with the side chain of Tyr 501. The aliphatic portion of the Arg side chain packs against the Tyr aromatic ring creating an enhanced hydrophobic contact in the double mutant compared to the N501Y single mutant (grey shading, Fig. 4A), while one hydrogen of the Arg guanidinium head group forms a hydrogen bond with the oxygen of the Tyr hydroxyl group (dashed lines, Fig. 4A). The geometries of the two residues are such that this pairing can be carried out without introducing any conformational strain, either in the backbone or in the side chains themselves. Thus, in the presence of N501Y, the mutation from Gln to Arg actually enhances binding rather than diminishing it.

As a control, we considered a second double mutant (L452R T478K) where no cooperativity is observed experimentally (Fig. 4C). The FEP calculations at 100ns predict values of both single and double mutant within 0.4 kcal/mol from the experiment and a cooperativity energy of −0.1 kcal/mol (same as SPR) (Fig. 4C). Thus, FEP accurately predicts the absence of cooperativity in the double mutant belonging to the Delta SARS-CoV-2 variant (Table S2).

## Discussion

We have carried out experimental and computational studies of a series of RBD mutants located in the RBD::ACE2 interface. The choice was based in part on their frequency in infective SARS-CoV-2 variants and in part because they have been extensively studied (20–63) and thus provided a useful data set to compare different approaches. A central goal has been to assess the ability of the FEP free energy perturbation methodology to predict the effect of mutations at the protein-protein interface on binding affinities but this in turn required an assessment of experimental accuracy. To this end we carried out SPR experiments on each mutant and compared our findings to those previously reported. We also applied easily accessible computational methods to our data set and compared their results to those obtained from FEP.

As shown in Table 1 and discussed above, there is excellent agreement between our SPR results and previous yeast display results from the Bloom group (20). Moreover, both sets of results are in the range reported in other studies. In particular, all methods agree on the identification of stabilizing mutations although the experimental values vary by as much as 1 kcal/mol, likely due to some of the issues discussed in Methods. As pointed out above, FEP yields the best correlation with our SPR results and, crucially, is the most effective in identifying stabilizing mutations (Table 2). In contrast, ML methods fail to yield meaningful correlations with experiment. This is not surprising since long simulations are often required to allow the system enough time to relax into a stable conformation while ML methods are based on data derived from fixed crystal structures. FoldX (a rigid backbone method) and Rosetta flex ddG (which accounts in part for backbone flexibility) are at least partially successful in identifying stabilizing mutations (Table 2).

In addition to the methods we were able to test on our data set, many other studies on RBD mutants have been reported. Table S4 lists published results on RDB::ACE2 on different sets of mutants than studied here but it is of interest to compare cases where there is overlap with previous work. Previous FEP studies (53, 55, 56) also carried out long simulations and all identify N501Y and N501T as stabilizing. TopNetTree(49), a topology-based ML method, does not perform well on our data set. The MM/GBSA study from the Bahar group (58) successfully predicts that N501Y is stabilizing but was not run on other mutants in our data set. Finally, Maranas and co-workers (47) used physical interactions extracted from MM/GBSA simulations to train a neural network to predict ΔΔG values (NN_MM-GBSA method). Since many of the mutants in our set were used in the NN_MM-GBSA training (Table S4) it is difficult to make comparisons with our own results. It is worth emphasizing that all simulation methods are at least partially successful in identifying stabilizing mutants thus, again, emphasizing the importance of allowing the system to relax.

Despite the success of the FEP approach revealed in this work, significant challenges remain. In previous work on a variety of systems, we have demonstrated that the FEP results correlate well with experiment (PCC values on the order of 0.6-0.8) and display RMS errors in the range expected for the OPLS4 (74) and related molecular mechanics force fields (0.5-1 kcal/mol) (8, 9, 14, 66–68). Precision beyond the above cited statistics is very difficult to obtain, indeed we have found (see Methods) that experimental reproducibility errors are typically on the order of 0.4 kcal/mol, even for the high quality SPR results that we report here.

As highlighted in this work, the major challenge in using FEP to predict mutation effects on protein-protein binding affinities, as opposed to small molecule binding, is the possibility of significant conformational change induced by mutation, for example if a buried charge is created by mutating a buried hydrophobic residue at the interface to one with a net charge. The key issue is whether the conformational changes required to make accurate predictions are accessible on the timescale of the FEP simulation. When the conformational changes consist primarily of side chain rearrangements and relatively minor backbone motion, FEP will typically deliver reliable results (as in N501Y). When there are significant backbone conformational changes, the accuracy of the FEP results will depend upon whether the barrier to conformational change can be surmounted on the time scale of the simulation.

In terms of overall accuracy, the results obtained here are consistent, in terms of both RMSD errors and correlation coefficients, with those we have reported previously in studying HIV derived gp120-antibody binding and also more diverse sets of protein-protein complexes via FEP simulations (8,9). The most significant errors may result from structural uncertainties (as in A475V) or from the prediction of overly unfavorable free energy changes (as in Q498R) which often result from electrostatic or steric clashes or from the burial of a charge in a hydrophobic pocket (9). However, mutations that are predicted to be highly destabilizing would be of little or no consequence in the context of binding optimization project since such mutations would be rejected in an initial screen even if converged results were obtained.

Our results suggest how FEP might be used in a strategy to optimize the binding of two proteins, for example in the optimization of antibodies. FEP calculations appear to have sufficient rank ordering capability to enable prioritization of specific single residue mutations. The initial step in a design strategy would be an exhaustive screening of single mutations at the various positions across the protein-protein interface (perhaps a few hundred to a few thousand calculations, which would require relatively modest expenditure, given the steadily decreasing cost of GPU-based computation). A key advantage of FEP is that analysis of trajectories can reveal insights as to the response of the interface to different perturbations, enhanced when needed by loop modeling procedures that more efficiently sample conformational space. These insights could then be translated into the investigation of a selected subset of double and triple mutations, with the goal of achieving favorable nonadditive effects. The understanding we gained about the Q498R N501Y double mutant could not have been accomplished by any of the other methods tested here.

It is important to clarify that, despite identifying cooperativity of the N501Y Q498R double mutant, the FEP results have not reached the point where the calculated values have experimental accuracy. It is then interesting to consider how a researcher interested in designing a stabilizing mutation would respond to calculated single mutant values of −1.2 kcal/mol for N501Y, +2.7 kcal/mol for Q498R and −1.4 kcal/mol for the double mutant. We would argue that the large calculated cooperativity energy of −2.9 kcal/mol, and the physical basis of this effect revealed by the simulations, would provide a strong hint that the double mutant is worth testing experimentally. Moreover, the experience we gained in this study would make that decision more likely. In conclusion, a careful exploration of a particular system of biological importance as enabled by FEP simulations would then appear to offer a way forward in many practical applications.

## Materials and Methods

### Dataset

We focused on missense RBD mutations at the interface with ACE2 that occurred most frequently in the US at the beginning of 2021 or were a part of known variants of concern. Among single point mutations with a frequency above 100 as of Jan 4, 2021, only seven mutations (S477N, N439K, N501Y, L452R, Y453F, S477R and S477I) were both missense and interfacial. To expand the dataset, we added missense interfacial mutations with a lower frequency (in the range of 10-100) if mutations were stabilizing or destabilizing in the study of Bloom and co-workers (20) while nearly neutral positions were ignored. Stabilizing mutations were of interest as potentially increasing infectivity of the virus, while destabilizing mutations were a necessary addition to create a balanced dataset for a proper testing of ΔΔG predictors. The K417T and Q498R mutations were added due to occurrence in the SARS-CoV-2 variants of concern and not based on frequency counts. The K417T mutation was added due to its emergence in Brazil (as a part of the Gamma variant) at the time. The Q498R mutant was studied in the context of double mutant effects alongside N501Y due to the co-occurrence of these two mutations in Omicron variant (Table S2) and a recent study on *in vitro* evolution suggesting cooperativity between the two mutations (64). Although the cumulative frequency (i.e. a number of times a given mutation has been found in the sequenced SARS-CoV-2 genomes) has changed during 2021, the majority of the most frequently observed mutations in January 2021 were still among the most recurrent mutations at the end of the year, with our data set including 16 out of 20 interfacial missense RBD mutations that had the highest frequency (>1000) on December 31, 2021 (Table S2).

Frequencies of SARS-CoV-2 mutations were obtained from the Mutation Tracker resource (https://users.math.msu.edu/users/weig/SARS-CoV-2_Mutation_Tracker.html) (75) that relies on data from the GISAID database of coronavirus genomes (https://www.gisaid.org/).

### ΔΔG calculations

All calculations presented in Table S3 were performed using as input a crystal structure of the SARS-CoV-2 RBD complex with ACE2 of highest available resolution (2.45Å, PDBID: 6m0j).

Mutabind2 predictions were run on the https://lilab.jysw.suda.edu.cn/research/mutabind2/webserver. mCSM-PPI2 predictions were submitted as a query to the following webserver: http://biosig.unimelb.edu.au/mcsm_ppi2/.

Standalone version (http://compbio.clemson.edu/saambe_webserver/standaloneCode.zip) was used for binding affinity calculations of SAAMBE-3D. FoldX calculations were performed as described in Sergeeva et al. (70). Rosetta flex ddG calculations were run using a standalone version (https://github.com/Kortemme-Lab/flex_ddG_tutorial) with the following parameters (considered to give optimized performance of this method (7)): nstruct = 35, max_minimization_iter = 5000, abs_score_convergence_thresh = 1.0, number_backrub_trials = 35000, backrub_trajectory_stride = 35000.

We used Schrödinger software (2021-2 release) and default FEP+ protocols (implementing guidelines for protein-protein interactions published earlier (8, 9)) for our predictions of binding affinity changes in the ACE2/RBD complex upon RBD mutations. The release incorporates a recently developed OPLS4 force field (74), replica exchange with solute tempering (REST) enhanced sampling methodology for mutated residues, and an improved grand canonical Monte Carlo (GCMC) protocol for sampling solvent molecules around mutated residues (76). FEP+ is a fully physics-based model that uses explicitly represented water. During FEP+ simulations, an alchemical transformation of a wild type amino acid residue into a mutant residue is conducted, which is implemented by running a series of separate molecular dynamics simulations (“lambda windows”) with varied energy weighting. The differences between each adjacent lambda window are first calculated using a perturbative expansion and then summed up to estimate the total free energy change between a wild-type and mutant states. To enhance the convergence of the free energy calculations, our default protocols use 12 lambda windows for charge conserving mutations and 24 lambda windows for the charge changing mutations. All mutations were run for 10 ns and 100 ns (see Table S4). Calculations of double mutant effects were performed by simultaneous alchemical transformation of the two mutated residues. This procedure minimizes errors associated with a more common FEP protocol where mutations are introduced sequentially. For example, the ΔΔG (Q498R N501Y) is predicted to be −1.4 kcal/mol using the simultaneous protocol and −0.6 kcal/mol using the sequential protocol. Of note, the results from the simultaneous protocol are in better agreement with the experimental value of −1.2 kcal/mol.

FEP+ requires GPU computing and the time required per single point mutation depends on the length of simulation (in nanoseconds), system size (in atoms), and type of mutation (with charge-changing mutations taking longer to compute compared to charge-neutral mutations as the number of lambda windows is twice as large). In the ACE2/RBD system (~100,000 atoms including explicit waters in the solvent box), the shortest 10 ns charge-neutral mutations take less than a day, while the longest 100 ns charge-changing simulations take ~2 weeks when complex and solvent legs are run in parallel with each simulation leg using 4GPUs.

### Protein Expression and Purification

The SARS-CoV-2 RBD wild-type and its mutants (residues 331-528), were cloned into the pVRC-8400 mammalian expression plasmid, with a C-terminal 6XHis-tag and an intervening HRV-3C protease cleavage site. Plasmid constructs were transfected into HEK293 cells using polyethyleneimine (Polysciences). Cell growths were harvested four days after transfection, and the secreted proteins were purified from supernatant by nickel affinity chromatography using Ni-NTA IMAC Sepharose 6 Fast Flow resin (Cytiva) followed by size exclusion chromatography on a Superdex 200 column (Cytiva) in 10 mM Tris, 150 mM NaCl, pH 7.4.

A plasmid encoding ACE2 residues 1-615 (pcDNA3-sACE2(WT)-8his) was a gift from Erik Procko (Addgene plasmid # 149268; http://n2t.net/addgene:149268;RRID:Addgene_149268) (77). This plasmid was then mutated to encode ACE2 residues 1-620, followed by a C terminal HRV-3C protease cleavage site, and an 8X HIS tag. This construct was transfected into ExpiHEK293 cells using Expifectamine according to manufacturer’s instructions. Supernatants were harvested seven days after transfection, and ACE2 was purified by nickel affinity chromatography using His60 Ni Superflow Resin (Takara) followed by size exclusion chromatography on a Superdex 200 column (Cytiva) in 10 mM Tris, 150 mM NaCl, pH 8.0. 400 ug of purified ACE2 was then digested overnight at 4 degrees with 20 units of HRV-3C protease (Thermo Scientific). Digested ACE2 was then incubated with His60 Ni Superflow Resin, which was then washed with 2 column volumes of 10 mM Tris, 150 mM NaCl, 5 mM imidazole, pH 8.0. The flow through and wash were determined by SDS-PAGE to contain cleaved ACE2, which was purified by size exclusion chromatography on a Superdex 200 column (Cytiva) in 10 mM Tris, 150 mM NaCl, pH 8.0.

### Surface Plasmon Resonance

SPR binding assays for monomeric ACE2 binding to RBDs were performed using a Biacore T200 biosensor, equipped with a Series S CM5 chip, at 25°C, in a running buffer of 10mM HEPES pH 7.4, 150mM NaCl, 0.1mg/mL BSA and 0.01% (v/v) Tween-20 at 25°C. Each RBD was captured through its C-terminal his-tag over an anti-his antibody surface, generated using the His-capture kit (Cytiva, MA) according to the instructions of the manufacturer.

During a binding cycle, each RBDs was captured over individual flow cells at approximately 250 RU. An anti-his antibody surface was used as a reference flow cell to remove bulk shift changes from the binding signals. Monomeric ACE2 was prepared at six concentrations in running buffer using a three-fold dilution series, ranging from 1.1-270 nM. Samples were tested in order of increasing protein concentration, with each series tested in triplicate. Blank buffer cycles were performed by injecting running buffer instead of the analyte, after two ACE2 injections to remove systematic noise from the binding signal. The association and dissociation rates were each monitored for 180s and 300s respectively, at 50μL/min. Bound RBD/Fab complexes were removed using a 10s pulse of 15 mM H_3_PO_4_ at 100μL/min, thus regenerating the anti-his surface for a new cycle of recapturing of each RBD, followed by a 60s buffer wash at 100μL/min. The data was processed and fit to 1:1 interaction model using the Scrubber 2.0 (BioLogic Software). The number in brackets for each kinetic parameter represents the error of the fit.

We have developed an SPR assay to measure the binding kinetics and affinities of interactions between ACE2 and wild-type or mutant RBD with the RBD captured to the chip to avoid compromised binding activity resulting from chemical immobilization or repeated surface harsh regeneration steps during the experiments. The SARS-CoV-2 RBD is a basic molecule with a pI ~9, so capture to the chip surface will also minimize artifacts such as non-specific interactions between the positively charged RBDs and the negatively charged dextran layer of the sensor chips at physiological pH. Monomeric ACE2 was flown over as analyte to avoid avidity effects resulting from using dimeric ACE2. Studies that have performed such experiments in both orientations (RBD tethered to the surface vs in solution as an analyte) showed that using RBD as an analyte yielded affinities that were approximately three-fold stronger for the wild type RBD/ACE2 interaction, suggesting the presence of such non-specific interactions (24). In our SPR experiments we have determined the K_D_ for wild type RBD binding to monomeric ACE2 is 162.9 nM (Fig. S1), consistent with similar K_D_s reported from other groups that have used similar methodologies to perform their biosensor-based measurements (21, 78, 79). Fig. S1A shows the binding kinetics for interactions of mutant RBD proteins with ACE2 and Fig. S1B shows the kinetic parameters along with affinities calculated for each binding interaction, while Table 1 and Fig. 4B list experimental changes in binding affinities (ΔΔG=RTln(K_D(MT)_/K_D(WT)_)) when RBD is mutated. Experimental reproducibility errors in our SPR data is expected not to exceed 15% according to previous estimates based on multiple independent measurements (70); this corresponds to ~0.4 kcal/mol experimental error in the ΔΔG(SPR) values reported in this study. Of 23 single point RBD mutations probed, four mutations were identified as stabilizing: N501Y, Y453F, S477N and N501T (Table 1), which could bind monomeric ACE2 with affinities stronger than the wild-type interaction with ACE2.

### Differences in Experimental Setup Affecting Binding Affinity Changes

Previously reported experimental binding affinity changes upon RBD mutation of the ACE2/RBD complex span different choice of protein constructs and orientation of molecules used in the binding assays.

The differences in the constructs lie in the choice of monomeric vs. multimeric forms of interacting proteins and selection of protein domain boundaries. Some studies relied on monomeric ACE2 and RBD (21, 22, 24, 25, 27–30), whereas others used at least one of the molecule in a multimeric form (trimeric spike, dimeric ACE2 or monomeric ACE2 fused to a dimeric Fc tag) (20, 25, 26, 31, 32, 64). RBD domain starts at residue 333 and ends at residue 526. It is common that constructs used in studies flank the RBD construct with a few residues before and after the domain boundary (e.g. 331-528 (this study), 331-531 (20), 328-531 (21, 22)) though some studies use constructs where such flanking regions are too long so they could result in non-specific binding (especially when containing unpaired cysteine residues, e.g. RBD 319-591 (25)). Poor selection of protein domain boundaries can affect protein folding/integrity when a construct has incomplete domain sequence (e.g. RBD 343-532 construct is missing a β-strand (23)).

Orientation of molecules in the binding experiments (which molecule is tethered to the chip in SPR) can affect both absolute and relative binding affinities (discussed in SPR methods). Studies using a highly positively charged RBD molecule as analyte and ACE2 immobilized on a chip (27–32) can be affected by non-specific binding of RBD to the chip. For example, using RBD as analyte in experiments measuring binding affinity of Alpha, Beta, Gamma, and Delta variants (80) results in stronger binding (by 0.4-0.6 kcal/mol) compared to a setup minimizing non-specific binding by immobilizing RBD on a chip (81). We used the latter as a reference to assess performance of ΔΔG on predicting double mutant effects of the Delta variant (Fig. 4C).

## Data Availability

All study data are included in the article and/or supporting information.

## ACKNOWLEDGMENTS

The work was supported by Melissa and Bill Gates Foundation, INV-016167 (to B.H., L.S., and R. F. We thank Peter Kwong and his lab for providing the monomeric ACE2 protein used in a subset of our experiments.

