## Supplemental figures and tables for "Free energy perturbation calculations of mutation effects on SARS-CoV-2 RBD::ACE2 binding affinity"

**This PDF file includes:**

Figures S1 to S3
Tables S1 to S4

A

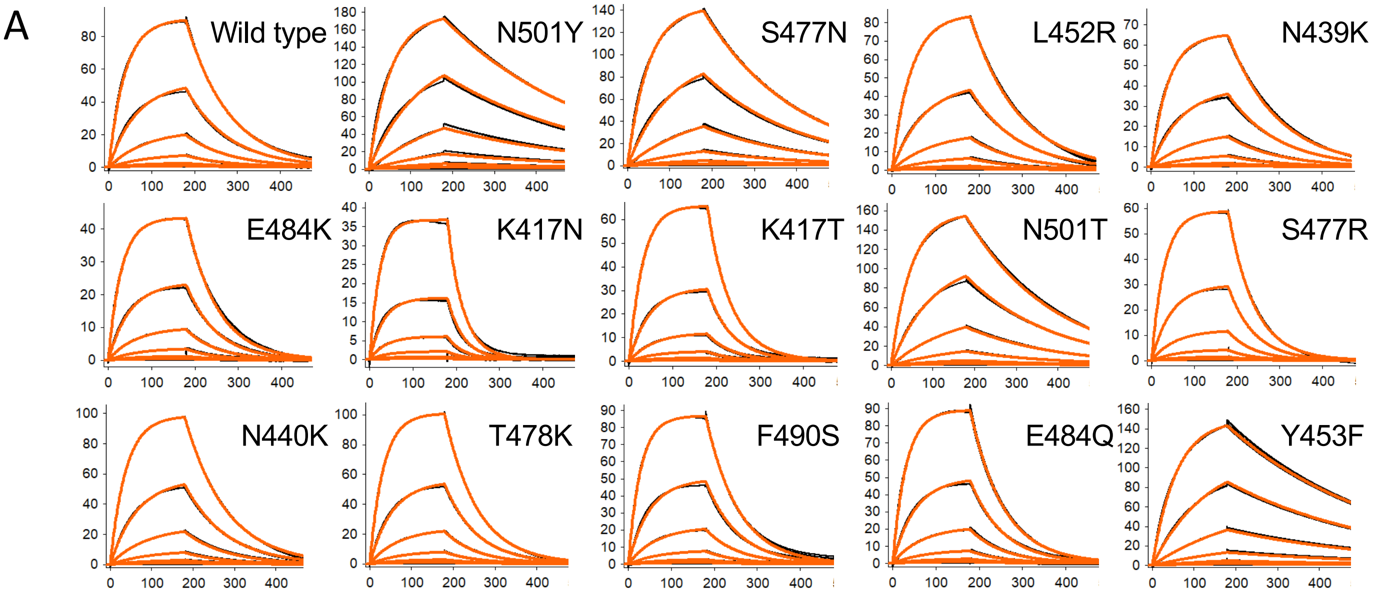

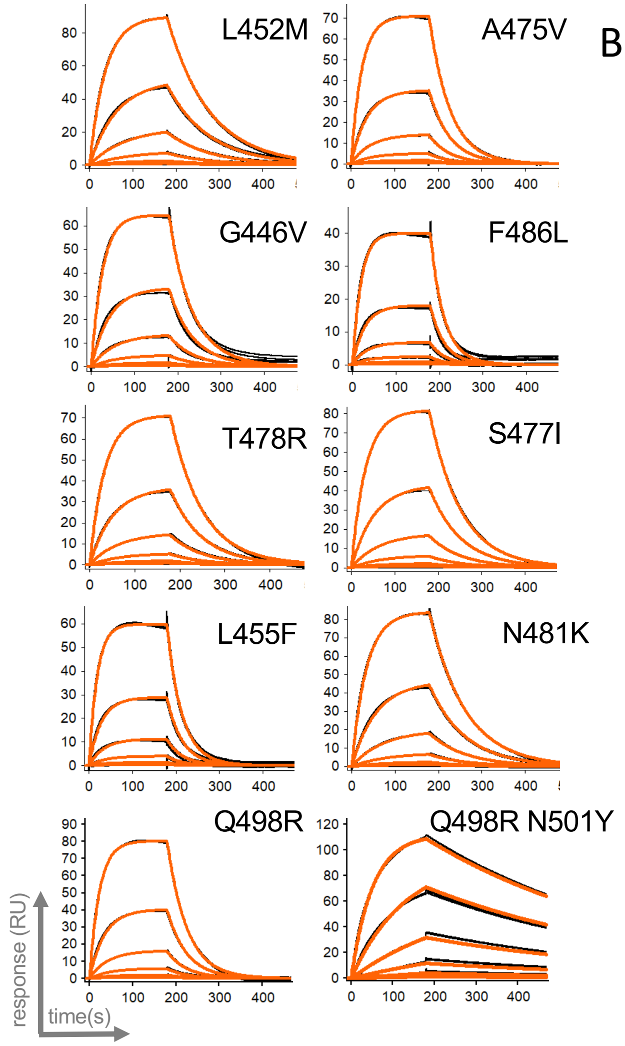

B

| RBD mutant | $k_a$ ($M^{-1}s^{-1}$) | $k_d$ ($s^{-1}$) | $K_D$ (nM) |
|---|---|---|---|
| RBD wild type | $6.18 \times 10^4$ | $1.01 \times 10^{-2}$ | 162.9 |
| RBD N501Y | $6.26 \times 10^4$ | $2.85 \times 10^{-3}$ | 45.6 |
| RBD S477N | $6.05 \times 10^4$ | $4.54 \times 10^{-3}$ | 75.2 |
| RBD L452R | $5.20 \times 10^4$ | $8.97 \times 10^{-3}$ | 172.5 |
| RBD N439K | $6.20 \times 10^4$ | $8.50 \times 10^{-3}$ | 137.0 |
| RBD E484K | $7.11 \times 10^4$ | $1.40 \times 10^{-2}$ | 196.4 |
| RBD K417N | $8.54 \times 10^4$ | $3.98 \times 10^{-2}$ | 466.0 |
| RBD K417T | $7.21 \times 10^4$ | $2.46 \times 10^{-2}$ | 342.0 |
| RBD N501T | $6.23 \times 10^4$ | $4.73 \times 10^{-3}$ | 76.0 |
| RBD S477R | $6.31 \times 10^4$ | $1.63 \times 10^{-2}$ | 257.7 |
| RBD N440K | $6.23 \times 10^4$ | $9.93 \times 10^{-3}$ | 159.3 |
| RBD T478K | $6.83 \times 10^4$ | $1.28 \times 10^{-2}$ | 187.6 |
| RBD F490S | $8.25 \times 10^4$ | $1.33 \times 10^{-2}$ | 160.5 |
| RBD E484Q | $7.30 \times 10^4$ | $1.31 \times 10^{-2}$ | 179.3 |
| RBD Y453F | $5.54 \times 10^4$ | $2.71 \times 10^{-3}$ | 48.9 |
| RBD L452M | $6.40 \times 10^4$ | $1.03 \times 10^{-2}$ | 161.3 |
| RBD A475V | $7.67 \times 10^4$ | $2.09 \times 10^{-2}$ | 273.0 |
| RBD G446V | $1.05 \times 10^5$ | $2.44 \times 10^{-2}$ | 233.6 |
| RBD F486L | $1.15 \times 10^5$ | $4.98 \times 10^{-2}$ | 430.0 |
| RBD T478R | $6.08 \times 10^4$ | $1.42 \times 10^{-2}$ | 234.2 |
| RBD S477I | $6.20 \times 10^4$ | $1.37 \times 10^{-2}$ | 221.8 |
| RBD L455F | $1.13 \times 10^5$ | $3.56 \times 10^{-2}$ | 315.7 |
| RBD N481K | $6.50 \times 10^4$ | $1.22 \times 10^{-2}$ | 187.6 |
| RBD Q498R | $1.02 \times 10^5$ | $2.20 \times 10^{-2}$ | 217.2 |
| Q498R N501Y | $8.15 \times 10^4$ | $1.84 \times 10^{-3}$ | 22.6 |

**Figure S1: SPR analysis of wt and mutant RBD/ACE2 binding affinities and kinetic parameters.** (A) SPR binding profiles for ACE2 binding to RBD and its mutants. The black traces represent experimental data, overlaid with the red traces which represent the 1:1 interaction model used to extrapolate kinetic parameters. (B) Kinetic parameters determined from the fits shown in A. The error of the fit for $k_a$ is (0.003-0.05)x$10^4$ $M^{-1}s^{-1}$, for $k_d$ is (0.004-0.6)x$10^{-3}$ $s^{-1}$ and for $K_D$ is 0.07-7 nM.

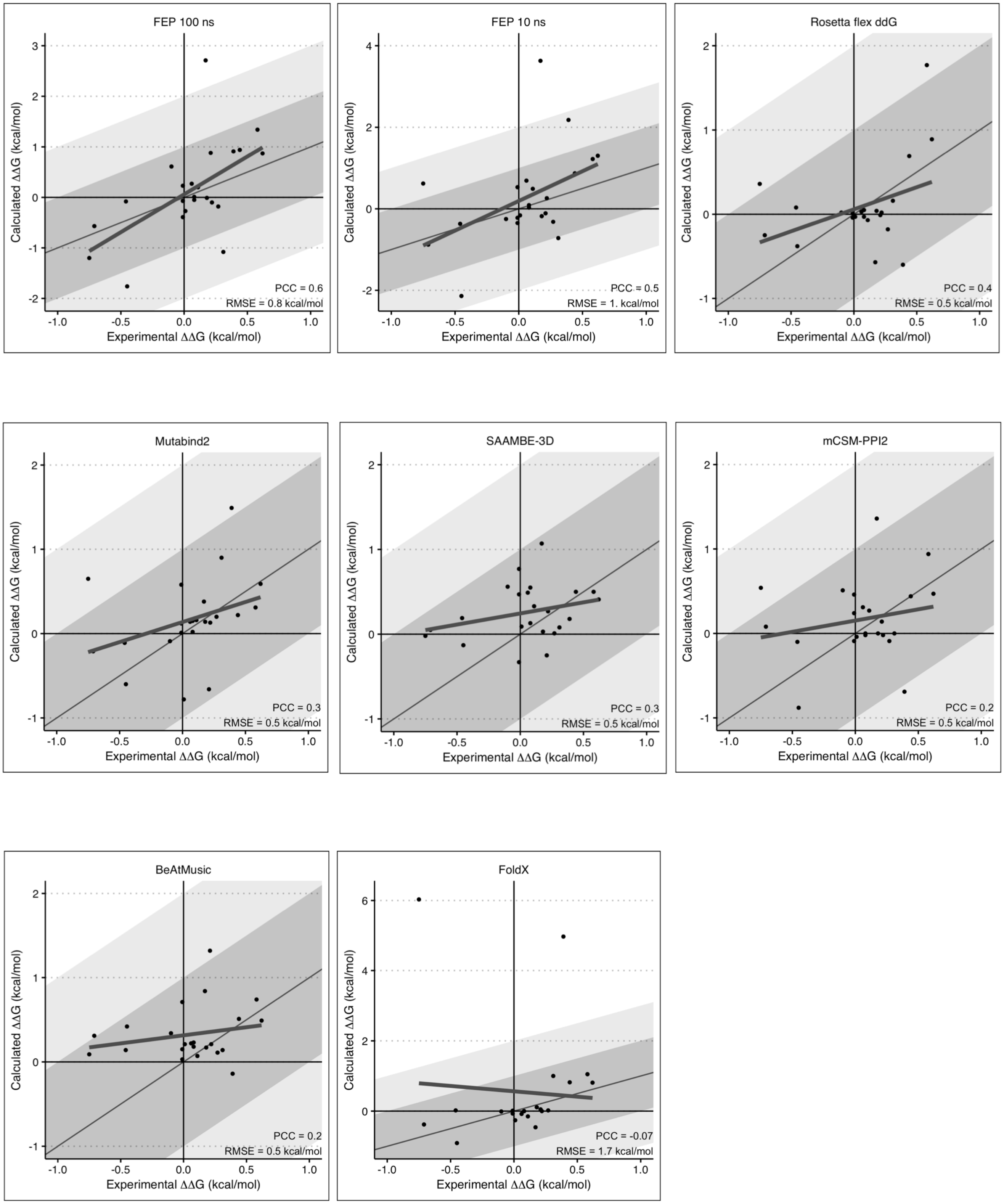

**Figure S2. Correlation plots for data in Table 2 and Table S3.**

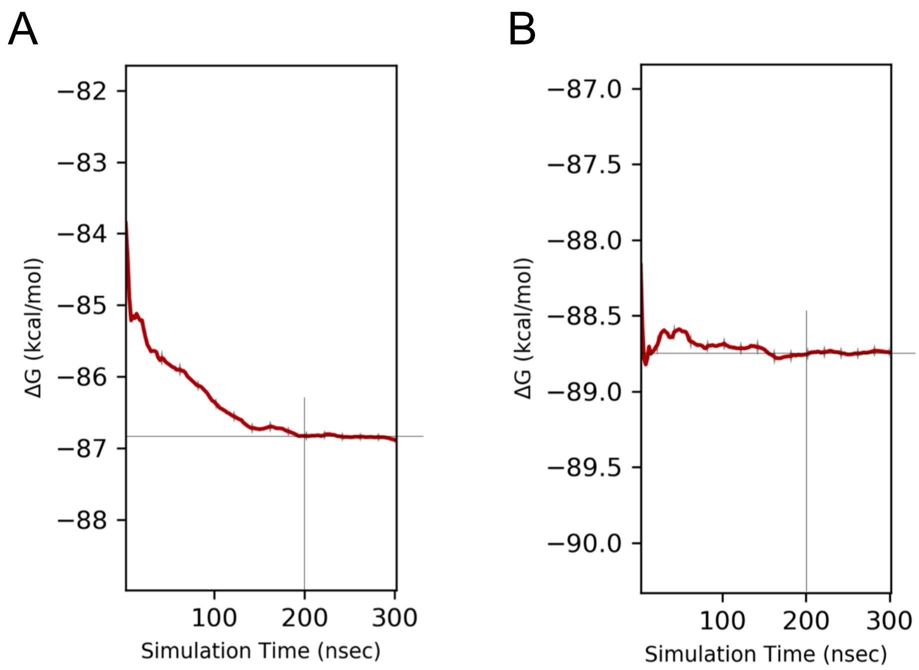

**Figure S3. Energy convergence plots for Q498R mutation.** Convergence plots of the FEP calculations for the complex leg (A) and the solvent leg (B) of the Q498R mutation simulated for 300ns to demonstrate that energy convergence is achieved at 200ns. ΔΔG (mutation) = ΔG (complex leg) – ΔG (solvent leg); ΔG = G(mutant) - G(wild-type).

### Table S1. Occurrence frequency of single point SARS-CoV-2 RBD mutations in the US in 2021.

| RBD mutation | Total frequency, Jan 04'21 | Total frequency, Jan 20'21 | Total frequency, Feb 19'21 | Total frequency, Mar 10'21 | Total frequency, April 18'21 | Total frequency, May 24'21 | Total frequency, June 7'21 | Total frequency, July 26'21 | Total frequency, Aug 31'21 | Total frequency, Oct 31'21 | Total frequency, Dec 12'21 | Total frequency, Dec 31'21 |
|---|---|---|---|---|---|---|---|---|---|---|---|---|
| L452R | 213 | 465 | 1141 | 1247 | 10056 | 25430 | 77577 | 180087 | 492276 | 1094585 | 1663119 | 1875810 |
| T478K | 13 | 35 | 177 | 190 | 4948 | 14146 | 62325 | 161860 | 467835 | 1069045 | 1637282 | 1849188 |
| N501Y | 373 | 469 | 11873 | 12081 | 169174 | 388294 | 614750 | 711234 | 778190 | 822805 | 837129 | 840571 |
| E484K | 82 | 192 | 487 | 592 | 9432 | 31142 | 60178 | 77492 | 97264 | 116482 | 122751 | 124058 |
| K417T | 0 | 0 | 17 | 40 | 1508 | 8481 | 23113 | 33295 | 47315 | 62212 | 65836 | 66643 |
| S477N | 12833 | 14448 | 16695 | 17160 | 22109 | 26139 | 29491 | 31840 | 33170 | 34071 | 34939 | 35592 |
| N439K | 3971 | 5435 | 6625 | 6872 | 10027 | 11899 | 14295 | 15617 | 16505 | 18474 | 19124 | 19388 |
| K417N | 19 | 21 | 90 | 102 | 1653 | 3993 | 6461 | 7733 | 9415 | 11179 | 11855 | 12188 |
| E484Q | 19 | 21 | 34 | 35 | 201 | 1062 | 2682 | 3102 | 3624 | 4953 | 7010 | 7926 |
| F490S | 25 | 27 | 37 | 42 | 388 | 1633 | 2740 | 3155 | 5971 | 6350 | 6572 | 6702 |
| G446V | 36 | 47 | 57 | 58 | 122 | 182 | 376 | 693 | 1272 | 2819 | 4179 | 4997 |
| N440K | 32 | 46 | 80 | 84 | 464 | 892 | 1741 | 2987 | 4091 | 4485 | 4847 | 4995 |
| S477I | 112 | 119 | 135 | 135 | 186 | 286 | 401 | 543 | 1012 | 1968 | 2982 | 3381 |
| N501T | 87 | 232 | 451 | 479 | 1382 | 1672 | 1880 | 2013 | 2265 | 2346 | 2429 | 2527 |
| A475V | 27 | 50 | 65 | 67 | 123 | 185 | 559 | 875 | 1382 | 1475 | 1560 | 1609 |
| L455F | 32 | 35 | 36 | 36 | 131 | 304 | 462 | 574 | 772 | 1063 | 1283 | 1373 |
| S477R | 144 | 250 | 321 | 329 | 538 | 740 | 812 | 839 | 847 | 883 | 907 | 924 |
| L452M | 54 | 62 | 66 | 67 | 137 | 382 | 632 | 713 | 791 | 854 | 874 | 896 |
| Y453F | 161 | 163 | 177 | 178 | 185 | 232 | 241 | 261 | 276 | 286 | 310 | 330 |
| T478R | 13 | 17 | 33 | 37 | 79 | 90 | 105 | 150 | 171 | 236 | 280 | 303 |
| Q498R | 0 | 0 | 0 | 0 | 0 | 0 | 0 | 5 | 12 | 35 | 71 | 207 |
| F486L | 90 | 90 | 138 | 138 | 139 | 142 | 144 | 148 | 150 | 153 | 155 | 156 |
| N481K | 13 | 37 | 47 | 47 | 65 | 98 | 95 | 103 | 109 | 129 | 134 | 134 |

### Table S2. RBD mutations present in select SARS-CoV-2 variants.

| Variant, WHO classification | Importance | PANGO Lineage | Country first detected | RBD Mutations |
|---|---|---|---|---|
| Alpha (α) | VOC | B.1.1.7 | United Kingdom | N501Y |
| Beta (β) | VOC | B.1.351 | South Africa | K417N, E484K, N501Y |
| Gamma (γ) | VOC | P.1 | Brazil | K417T, E484K, N501Y |
| Delta (δ) | VOC | B.1.617.2 | India | L452R, T478K |
| Epsilon (ε) | VOI->VUM->reclassified | B.1.427/B.1.429 | USA | L452R |
| Zeta (ζ) | VOI->VUM->reclassified | P.2 | Brazil | E484K |
| Eta (η) | VOI->VUM | B.1.525 | Nigeria | E484K |
| Theta (θ) | VOI->VUM->reclassified | P.3 | The Philippines | E484K, N501Y |
| Iota (ι) | VOI->VUM | B.1.526 | USA | S477N, E484K |
| Kappa (κ) | VOI->VUM | B.1.617.1 | India | L452R, E484Q |
| Lambda (λ) | VOI | C.37 | Peru | L452Q, F490S |
| Mu (μ) | VOI | B.1.621 | Colombia | R346K, E484K, N501Y |
| Omicron (o) | VOC | B.1.1.529 | South Africa | G339D, S371L, S373P, S375F, K417N, N440K, G446S, S477N, T478K, E484A, Q493R, G496S, Q498R, N501Y, Y505H |
| "Stealth" omicron | VOC | BA.2 | South Africa | G339D, S371F, S373P, S375F, T376A, D405N, R408S, K417N, N440K, S477N, T478K, E484A, Q493R, Q498R, N501Y, Y505H |
| - | - | B.1.258 | Scotland | N439K |
| - | - | A.28 | France | E484K, N501T |
| - | - | B.1.1.298 (cluster 5) | Denmark/Netherlands (mink) | Y453F |

Single point mutations experimentally probed in the current study are highlighted in blue.

**Importance**

| | |
|---|---|
| VOC | variant of concern |
| VOI | variant of interest |
| VUM | variant under monitoring |
| reclassified | Former VOCs/VOIs/VUMs, including their descendent lineages, that have been reclassified based on at least one the following criteria:<br>(1) the variant is no longer circulating at levels of global public health significance,<br>(2) the variant has been circulating for a long time without any impact on the overall epidemiological situation, or<br>(3) scientific evidence demonstrates that the variant is not associated with any concerning properties. |

**Table S3. Comparison of FEP results at 10ns and 100ns to experimental SPR values.** Color code and notations as in Table 2.

| RBD mutation | ΔΔG, experiment, SPR, kcal/mol | ΔΔG FEP+, 10ns | ΔΔG FEP+, 100ns |
|---|---|---|---|
| N501Y | -0.8 | 0.6 | -1.2 |
| Y453F | -0.7 | -0.9 | -0.6 |
| S477N | -0.5 | -0.4 | -0.1 |
| N501T | -0.5 | -2.1 | -1.8 |
| N439K | -0.1 | -0.3 | 0.6 |
| N440K | 0.0 | -0.2 | -0.4 |
| F490S | 0.0 | -0.4 | -0.1 |
| L452M | 0.0 | 0.5 | 0.2 |
| L452R | 0.0 | -0.2 | -0.3 |
| E484Q | 0.1 | 0.7 | 0.3 |
| T478K | 0.1 | 0.0 | 0.0 |
| N481K | 0.1 | 0.1 | -0.1 |
| E484K | 0.1 | 0.5 | 0.2 |
| Q498R | 0.2 | 3.6 | 2.7 |
| S477I | 0.2 | -0.2 | 0.0 |
| G446V | 0.2 | -0.1 | 0.9 |
| T478R | 0.2 | 0.3 | -0.1 |
| S477R | 0.3 | -0.3 | -0.2 |
| A475V | 0.3 | -0.7 | -1.1 |
| L455F | 0.4 | 2.2 | 0.9 |
| K417T | 0.4 | 0.9 | 0.9 |
| F486L | 0.6 | 1.2 | 1.3 |
| K417N | 0.6 | 1.3 | 0.9 |
| | PCC | 0.5 | 0.6 |
| | RMSE | 1.0 | 0.8 |
| | PCC ϕ (stabilizing) | 0.6 | 0.6 |

**Table S4. Comparison of binding affinity changes measured and calculated in the current study to previously reported computational predictions.** ΔΔG values are given in units of kcal/mol.

| RBD mutation | ΔΔG, experiment, SPR, kcal/mol | ΔΔG FEP+, 100ns | ΔΔG FEP, Fratev | ΔΔG FEP, Huynh | ΔΔG FEP, Gumbart | ΔΔG TopNetTree, Wei | ΔΔG AMBER/MMPBSA, Pricl | ΔΔG PRODIGY/ MMGBSA, Wei | ΔΔG PRODIGY/ MMGBSA, Bahar | ΔΔG PRODIGY/ MMPBSA, Zonta | ΔΔG NN_MMGBSA, Maranas with * | *-mutations NN_MMGBSA was trained on |
|---|---|---|---|---|---|---|---|---|---|---|---|---|
| N501Y | -0.8 | -1.2 | -1.4 | -0.8 | -4.5 | 0.0 | | 0.1 | -0.4 | 0.4 | -0.1 | N501Y* |
| S477N | -0.5 | -0.1 | | | | -0.1 | | | | | -0.1 | S477N (S477D*) |
| L452R | 0.0 | -0.3 | | | | -2.5 | | | | | -0.1 | L452R (L452K,Q*) |
| N439K | -0.1 | 0.6 | | | | -1.4 | | | | | -0.1 | N439K* |
| E484K | 0.1 | 0.2 | | | -1.3 | -0.9 | | -0.3 | | -0.1 | -0.1 | E484K (E484R*) |
| K417N | 0.6 | 0.9 | 0.6 | | 0.4 | -1.8 | | | | 0.1 | 0.3 | K417N (K417E,R,V*) |
| K417T | 0.4 | 0.9 | | | | -1.1 | 1.5 | | | 0.2 | 0.3 | K417T (K417E,R,V*) |
| N501T | -0.5 | -1.8 | -1.9 | | | -1.4 | -0.3 | | | | -0.1 | N501T* |
| S477R | 0.3 | -0.2 | | | | -0.3 | | | | | | |
| N440K | 0.0 | -0.4 | | | | -0.4 | | | | | -0.1 | N440K* |
| T478K | 0.1 | 0.0 | | | | -0.2 | | | | | -0.1 | |
| F490S | 0.0 | -0.1 | | | | -0.6 | | | | | -0.1 | F490S (F490K*) |
| E484Q | 0.1 | 0.3 | | | | 1.2 | | | | | -0.1 | E484Q (E484R*) |
| Y453F | -0.7 | -0.6 | | | | -0.6 | | | | | -0.1 | Y453F* |
| L452M | 0.0 | 0.2 | | | | -2.4 | | | | | | |
| A475V | 0.3 | -1.1 | | | | 1.1 | | | | | | |
| G446V | 0.2 | 0.9 | | | | 0.4 | | | | | | |
| F486L | 0.6 | 1.3 | | | | -0.7 | | | | | | |
| T478R | 0.2 | -0.1 | | | | 0.1 | | | | | | |
| S477I | 0.2 | 0.0 | | | | -0.3 | | | | | | |
| L455F | 0.4 | 0.9 | | | | 0.7 | | | | | | |
| N481K | 0.1 | -0.1 | | | | 0.1 | | | | | | |
| Q498R | 0.2 | 2.7 | | | | -0.3 | | | | | | |

| | | | | | | | | | | | |
|---|---|---|---|---|---|---|---|---|---|---|---|
| | PCC | 0.6 | | | | 0.0 | | | | | 0.7 |
| | RMSE | 0.8 | | | | 1.1 | | | | | 0.3 |
